# The landscape of alternative polyadenylation in single cells of the developing mouse embryo

**DOI:** 10.1101/2021.01.21.427498

**Authors:** Vikram Agarwal, Sereno Lopez-Darwin, David R. Kelley, Jay Shendure

## Abstract

3′ untranslated regions (3′ UTRs) post-transcriptionally regulate mRNA stability, localization, and translation rate. While 3′-UTR isoforms have been globally quantified in limited cell types using bulk measurements, their differential usage among cell types during mammalian development remains poorly characterized. In this study, we examined a dataset comprising ∼2 million cells spanning E9.5–E13.5 of mouse embryonic development to quantify transcriptome-wide changes in alternative polyadenylation (APA). We observe a global lengthening of 3′ UTRs across embryonic stages in all cell types, although we detect shorter 3′ UTRs in hematopoietic lineages and longer 3′ UTRs in neuronal cell types within each stage. An analysis of RBP dynamics identifies ELAV-like family members, which are concomitantly induced in neuronal lineages and developmental stages experiencing 3′-UTR lengthening, as putative regulators of APA. By measuring 3′-UTR isoforms in an expansive single cell dataset, our work provides a transcriptome-wide and organism-wide map of the dynamic landscape of alternative polyadenylation during mammalian organogenesis.

## INTRODUCTION

During transcriptional elongation, the cleavage and polyadenylation machinery governs the specification of the 3′ terminal end of an mRNA^1^. This regulated process can generate a diversity of 3′-UTR isoforms for any given gene, dramatically altering the 3′-UTR length and sequence of the resulting mature transcripts^2^. This phenomenon, known as alternative polyadenylation (APA), has been observed in over 70% of mammalian genes^3,4^. Alternative 3′-UTR isoforms bind to different sets of microRNAs and RNA-binding proteins, which collectively modulate a multitude of post-transcriptional gene regulatory mechanisms^5^. These include changes in mRNA localization^6^, degradation rates^7–9^, and translational efficiency^10^. The differential abundance of a variety of nuclear factors serves to regulate APA in a cell-type-specific manner^1^. Abnormal regulation of the cleavage and polyadenylation machinery has also been associated with hyperproliferative or disease states such as cancer^11–13^.

Techniques to directly measure APA in the transcriptome largely rely upon the isolation of RNA from bulk tissue, resulting in an average readout of the landscape of 3′ ends in a heterogeneous population of cells. Existing 3′-end sequencing methods include 3′-seq/3SEQ^14,15^, 3P-seq^8,16^, PAS-seq^17^, 3′READS^18^, PolyA-seq^3^, and 2P-seq^19^. The successful application of these methodologies in mammalian cells has led to the annotation of hundreds of thousands of polyadenylation sites (PAS) in both human and mouse genomes^20,21^. Bulk 3′-end sequencing and similar transcriptomic data have guided the observation that 3′ UTRs generally lengthen during mammalian embryogenesis^22^, with proliferating cell types such as blood exhibiting shortened 3′ UTRs^11,12^ and neuronal ones exhibiting lengthened 3′ UTRs^17,23^.

In contrast to bulk methods, single-cell RNA sequencing (scRNA-seq) protocols capture a rich diversity of individual cell types, with many protocols enriching for mRNA 3′ ends via poly(A) priming^24–29^. Thus, these technologies inherently offer an unprecedented opportunity to observe APA events during the process of cellular differentiation. They also enable the decomposition of complex tissues into individual cell types, enabling the assessment of APA with greater cell type resolution. Although a proof-of-concept study has demonstrated the utility of scRNA-seq data in evaluating APA^30^, such methods have not yet been applied to investigate more comprehensive scRNA-seq datasets such as those capturing dozens of cell types during a mammalian developmental time course^31,32^. In this study, we examined APA using MOCA (“mammalian organogenesis cell atlas”), a dataset comprising single nucleus transcriptional profiling of ∼2 million cells encompassing 38 major cell types across five stages (*i.e.*, E9.5, E10.5, E11.5, E12.5, and E13.5) of mouse embryonic development^31^.

## RESULTS

### An integrated annotation set of 3′ UTRs and poly(A) sites to evaluate APA

Given the reliance of many scRNA-seq protocols on poly(A) priming, such methods enrich for both mRNA 3′ ends as well as internal A-rich stretches of homopolymers. Thus, internal priming artifacts obscure accurate quantitation of APA, even more so in datasets in which immature mRNAs (*i.e.*, without excised introns) are isolated from the nucleus, as is the case with the sci-RNA-seq3 protocol used in MOCA^31^. We therefore sought to develop a simple computational method to deconvolve the data to specifically isolate and quantify mRNA 3′ ends. Towards this goal, we built integrated databases of poly(A) site (PAS) and 3′-UTR annotations to guide the interpretation of which subset of mapped reads were supported by orthogonal evidence to reflect authentic 3′ termini, as opposed to A-rich sites internal to a mature or nascent transcript. In doing so, our goal was to minimize the shortcomings of any individual database, each of which utilize different data sources and strategies for PAS and 3′-UTR annotation.

To generate a reliable PAS set, we considered three of the most comprehensive mouse PAS annotation databases available with respect to the mm10 mouse genome build: Gencode M25^33^, which contains 56,592 PASs; PolyA_DB v3^20^, which contains 128,052 PASs; and PolyASite 2.0^21^, which contains 108,938 PASs. We intersected the PASs from each pair of these resources to evaluate the consistency among databases. While the majority of sites were present in at least two databases, 40.0%, 29.4%, and 30.4% were unique to PolyA_DB, PolyASite, and Gencode, respectively (**Fig. 1A**). To verify the reliability of PASs present in only a single database (and therefore the most likely to contain false positives), we plotted the profile of nucleotide frequencies in the ±50nt region surrounding the annotated cleavage and polyadenylation sites (**Supplementary Fig. 1A**). The unique PASs of each resource exhibited profiles consistent with positionally-enriched mammalian motifs known to guide mRNA cleavage, including several U-rich motifs, the upstream AAUAAA motif, and the downstream GU-rich motif^34^. Moreover, we detected a strong enrichment of reads mapping immediately upstream of this set of PASs, with the strongest enrichment spanning the -300nt to +20nt region around the PAS (**Supplementary Fig. 1B**). Given that each PAS database was enriched in known PAS motifs, associated with mapped reads, and held information complementary to the other databases, we carried forward an integrated PAS set derived from the union of the three databases. This integrated PAS set recapitulated these same characteristics, exhibiting both consistency with known PAS motifs and strong read enrichment upstream of the sites (**Fig. 1B,C**). Finally, to link PASs to specific genes, we utilized our previous 3′-UTR annotation pipeline (**Methods**)^9,35^ to establish an integrated set by carrying forward the longest 3′ UTRs from four resources: i) Gencode M25^33^, ii) RefSeq^36^, iii) 3′ UTRs with extreme lengthening^23^, and iv) bulk 3P-seq-based annotations derived from ten mouse tissues and cell lines^8^. This integrated 3′-UTR annotation set helped minimize the possibility that a PAS may be annotated outside of a known 3′ UTR and thus remain unlinked to a specific gene.

**Figure 1.**
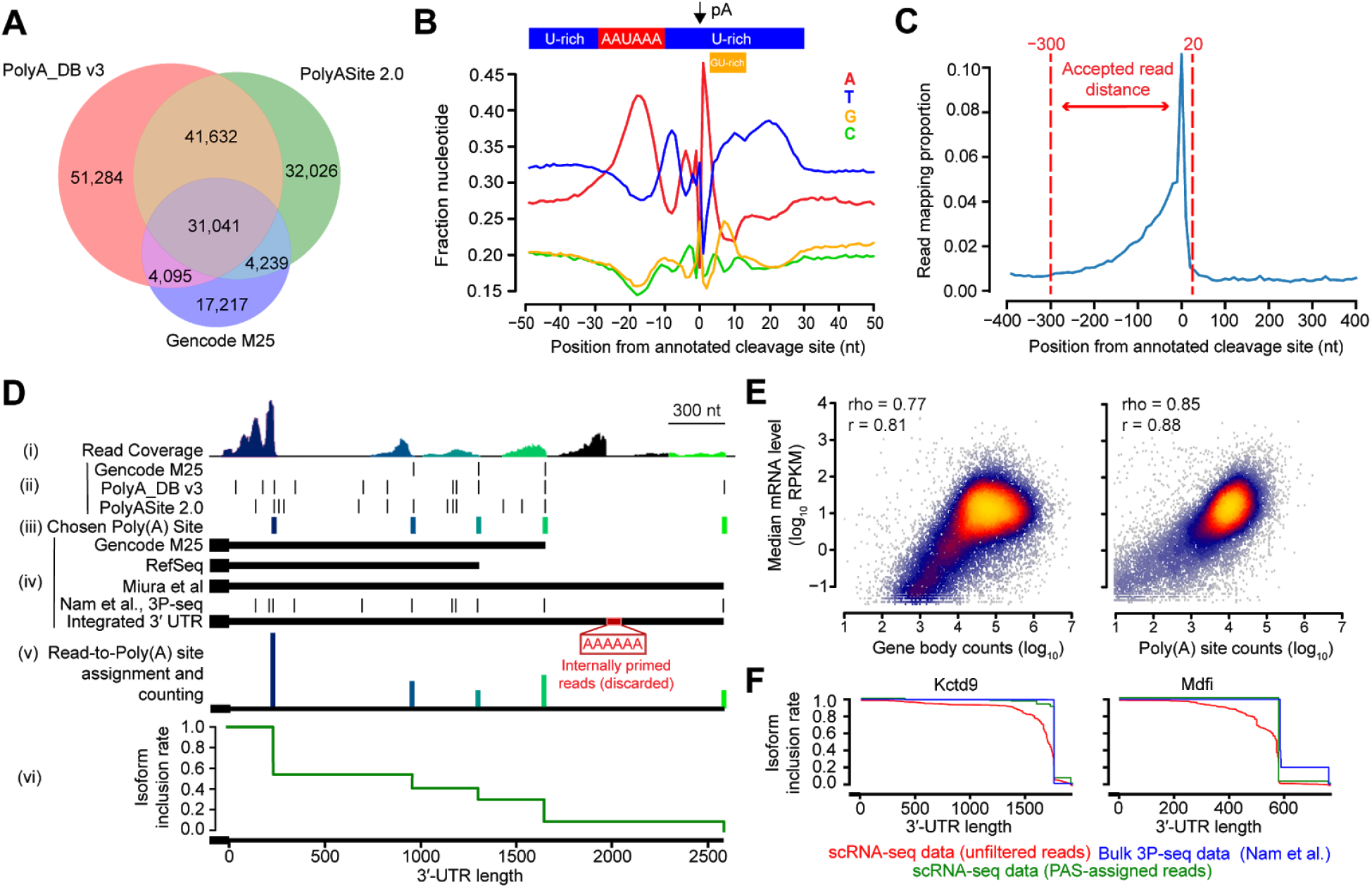
A computational pipeline to accurately quantify 3′-UTR isoform abundances from scRNA-seq data. **(A)** Venn diagram of a set of three PAS annotation resources and their degree of intersection. A PAS intersecting within ±20nt from another was considered an intersecting hit to account for the heterogeneity of the cleavage and polyadenylation machinery ^16^. **(B)** Profile of nucleotide frequencies in the ±50nt vicinity of the annotated cleavage site position, derived from the union of the three databases. Shown above the plot are the known positionally-enriched mammalian motifs known to guide mRNA cleavage^34^. **(C)** Distribution of scRNA-seq reads mapping within the ±400nt vicinity of the annotated cleavage site position, derived from the union of the three databases. To avoid an ambiguous signal, the analysis was restricted to PASs not within the same ±400nt window as another PAS. Data is binned at 5nt resolution. Shown within the dotted red lines are the acceptable distance thresholds to associate a read to an annotated PAS. See also **Supplementary Fig. 1** for comparisons of (B-C) for each individual PAS database. **(D)** Schematic depicting the association of each scRNA-seq read to a PAS in order to quantify relative PAS abundances for a gene. Shown from top to bottom are: (i) The read coverage of scRNA-seq reads mapped to the gene. (ii) The three PAS annotation resources considered, showing the location of each PAS along the 3′ UTR. (iii) The subset of chosen PASs to which reads were greedily assigned, colored from blue to green to indicate which reads from the coverage plot were assigned to them. (iv) The three gene annotation databases integrated with bulk 3P-seq data from ten tissues and cell lines^8^ to identify the longest known 3′ UTR. This integrated 3′ UTR was used to associate PASs to the gene. (v) A visualization of relative 3′-UTR isoform abundances after read-to-PAS assignment, with vertical lines at each chosen PAS proportional to the assigned number of read counts. Reads not overlapping within the -300 to +20 vicinity of a known PAS were treated as likely internal priming artifacts and discarded. (vi) The resulting isoform inclusion rate (IIR) plot to quantify the cumulative proportion of 3′-UTR isoforms remaining along the length of a 3′ UTR. See also **Supplementary Table 1** for the integrated 3′-UTR database and gene annotations. **(E)** Scatter plots comparing gene expression levels estimated using scRNA-seq read abundances mapping to the full gene body (left panel) or the sum of reads mapping to PASs (right panel), relative to median gene expression levels from bulk RNA-seq data^37^ (n = 19,517 protein-coding genes). Regions are colored according to the density of data from light blue (low density) to yellow (high density). Shown are the corresponding Pearson (r) and Spearman (rho) correlations for each comparison. See also **Supplementary Fig. 2** for sequence features explaining biased estimates in the gene body approach. **(F)** Shown are IIR plots for two genes, comparing the profiles for the raw scRNA-seq data and post-processed data after read-to-PAS assignment with respect to the profile for bulk 3P-seq data ^8^ as a gold standard. Slight vertical jitter was added for enhanced line visibility. See also **Supplementary Fig. 3** for a global comparison among all genes.

Using our integrated PAS and 3′-UTR databases (**Fig. 1D**), we sequentially filtered our scRNA-seq reads from MOCA to focus on the subset mapping to 3′ UTRs within the -300 to +20 vicinity of a known PAS (**Fig. 1C**). Due to the abundant mapping of reads to introns in the nucleus-derived MOCA dataset (**Supplementary Fig. 1C**), generally representing internal priming within unspliced transcripts, there was nearly a 10-fold loss in read counts after iterative steps of filtering; however, over 200 million reads were carried forward (**Supplementary Fig. 1D**). We then counted reads passing the filtering steps towards the single annotated PAS in its vicinity, enabling the tabulation of read counts associated with each PAS (**Fig. 1D**). In ambiguous cases in which a read was located in the vicinity of multiple PASs, we greedily assigned the read to count towards the PAS harboring the most uniquely assignable reads. Finally, based upon the relative counts assigned to each PAS for a given gene, we visualized the “isoform inclusion rate” (IIR), reflecting the proportion of 3′-UTR isoforms which include a given nucleotide position^8,9,35^ (**Fig. 1D**).

To validate that these filtering and read-to-PAS assignment procedures led to reliable results, we performed two quality control (QC) comparisons. As a first QC, reasoning that the removal of internal priming artifacts should improve the quantitation of relative gene expression levels, we compared the relationship between PAS counts and median gene expression levels computed across a panel of 254 mouse RNA-seq samples^37^. While the traditional method of counting reads in the gene body displayed a strong correlation to median expression levels (Pearson r = 0.81, Spearman rho = 0.77), it displayed a clear bias in inflating estimates for a large cohort of genes (**Fig. 1E**). Considering only our filtered PAS-assigned reads ameliorated this bias, which led to a stronger correlation to relative mRNA expression levels (Pearson r = 0.88, Spearman rho = 0.85) (**Fig. 1E**). We speculated that the bias in the gene body method relative to the PAS-assigned read counting method could be explained by the over-abundance of intron-mapping reads (**Supplementary Fig. 1C**) and enrichment of A-rich stretches that nucleate the production of internal priming artifacts. Indeed, a lasso regression model trained to predict the difference between the two strategies confirmed that intron length was strongly associated with inflated counts; moreover, “AAA” was the top-ranked of all 3-mers associated with inflated counts in the gene body (Pearson r = 0.56, Spearman rho = 0.58, **Supplementary Fig. 2**).

As a second QC, we evaluated the similarity between our IIR profiles to those derived from bulk 3P-seq data^8^. We considered the latter as a gold standard in accurately quantifying PAS abundances due to the involvement of a splint-ligation step in the 3P-seq protocol, which specifically removes internal priming artifacts^16^. We found that the IIR profiles for our PAS-assigned reads more strongly mirrored those of bulk data for two representative genes (**Fig. 1F**). Quantifying the deviation from bulk as the Mean Absolute Deviation (MAD) (**Supplementary Fig. 3A**) allowed us to measure the deviations between our pre- and post-processed data to bulk 3P-seq measurements. Applying this metric globally to all genes uncovered that 78% of genes exhibited improved similarity to bulk after the reads were assigned to PASs; moreover, 47% of genes achieved MAD <= 0.1 after read-to-PAS assignment, relative to only 7% of genes beforehand (**Supplementary Fig. 3B**). Inspection of IIR profiles for nine representative genes further confirmed the general improvement in consistency with bulk 3P-seq data (**Supplementary Fig. 3C**).

### Global differences in 3′-UTR length across mouse cell types and developmental time

Having assigned reads to PASs and linked them to genes, we next sought to evaluate global properties of 3′-UTR shortening and lengthening (*i.e.*, as quantified by differential PAS usage) across cell types and developmental time. Towards this goal, we computed a “gene by cell” sparse matrix of the mean length among all 3′-UTR isoforms, weighted by their respective counts. For each gene, we then computed each cell’s deviation from the mean of 3′-UTR lengths across cells, considering only non-missing values. Finally, for each cell, we computed the mean of these deviations across genes as a measure of the global behavior of the transcriptome through the perspective of APA. We projected these measurements onto a global map of 38 t-SNE clusters representing all major mouse cell types^31^. Highlighting the top ten ranked t-SNE clusters with the largest differences, we discovered the greatest average 3′-UTR lengths among stromal cells and three neuronal cell types; in contrast, the shortest lengths occurred in three blood cell types, hepatocytes, chondrocytes, and osteoblasts (**Fig. 2A**). Sub-clustering each of the 38 t-SNE clusters reinforced these findings, but revealed additional heterogeneity within each cell type (**Supplementary Fig. 4**). Segregating our dataset by the five sampled timepoints, we observed an apparent global 3′-UTR lengthening across developmental time (**Fig. 2B**). Finally, to quantify the joint impact of cell type and developmental stage, we computed the average behavior among genes and cells associated with each of 38 t-SNE clusters and 5 developmental stages. Partitioning the data in this manner reinforced our observation that average 3′-UTR length increased in nearly every cell type as developmental time progressed (**Fig. 2C**). Neuronal cell types clustered together and exhibited the greatest 3′-UTR lengthening relative to other clusters at E13.5; in contrast, blood cell types exhibited highly shortened 3′ UTRs at E9.5 and grew until E13.5 to mean lengths similar to those of other cell types at E9.5 (**Fig. 2C**).

**Figure 2.**
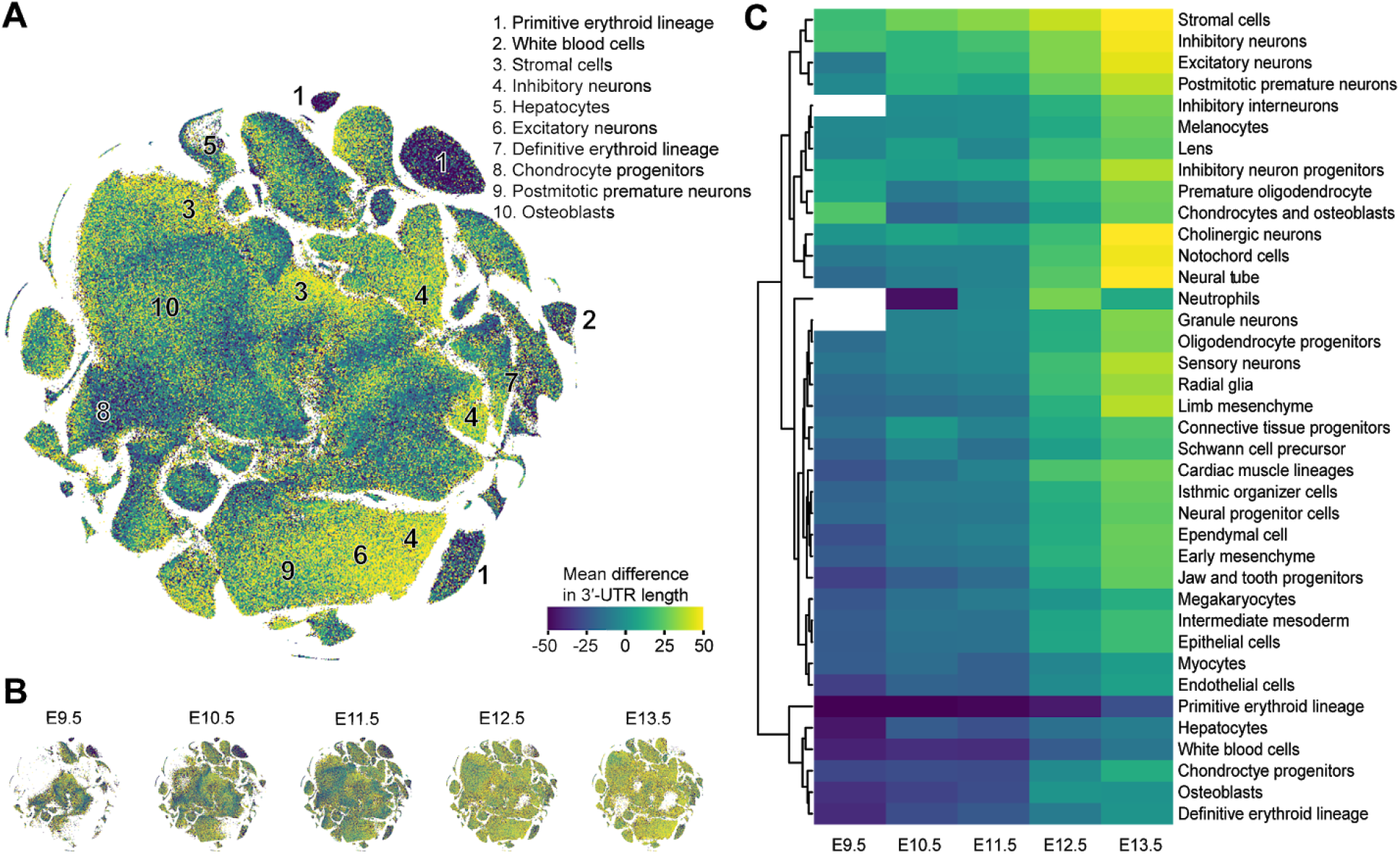
Differential 3′-UTR lengthening among diverse cell types and developmental stages. **(A)** t-SNE embedding of all cells from all developmental stages sampled^31^, with each cell colored according to the mean difference in 3′-UTR lengths across all genes. The top ten ranked clusters with the greatest differences are annotated according to their corresponding cell type. **(B)** Shown are the same embeddings and color scales as those in panel (A), except after partitioning the dataset into its five composite developmental stages (spanning E9.5-E13.5). **(C)** Heatmap of the mean difference in 3′-UTR length after aggregating cells from each developmental stage and cell type, derived from each of 38 t-SNE clusters. Color scales are the same as those shown in panel (A). Missing values (shown in white) correspond to instances with too few (<20) cells to accurately estimate. Heatmap is clustered by Euclidean distance as a distance metric. See also **Supplementary Fig. 4** for comparisons among t-SNE subclusters.

Next, we evaluated differences in global 3′-UTR length with respect to developmental trajectories computed using UMAP, an embedding that more faithfully recapitulates cell–cell relationships and intermediate states of differentiation relative to t-SNE. Evaluating ten developmental UMAP trajectories^31^, we again observed a global lengthening in 3′ UTRs in nearly every trajectory (**Fig. 3A**). Mirroring our previous findings, the neural tube/notochord and the neural crest trajectories (capturing neurons of the peripheral nervous system) showed the greatest lengths relative to other cell types at E13.5, while the hematopoiesis trajectory displayed the shortest lengths relative to other cell types at E9.5 (**Fig. 3A**). A visual comparison of these three trajectories with respect to changes in both developmental time and 3′-UTR length showed that the process of 3′-UTR lengthening occurred contemporaneously with cellular differentiation, with gradients of lengthening emerging in intermediate cellular states (**Fig. 3B-D**). Notably, in the hematopoiesis trajectory, a major difference in 3′-UTR length could be explained by the switch from primitive to definitive erythropoiesis, rather than gradual lengthening within either lineage (**Fig. 3D**).

**Figure 3.**
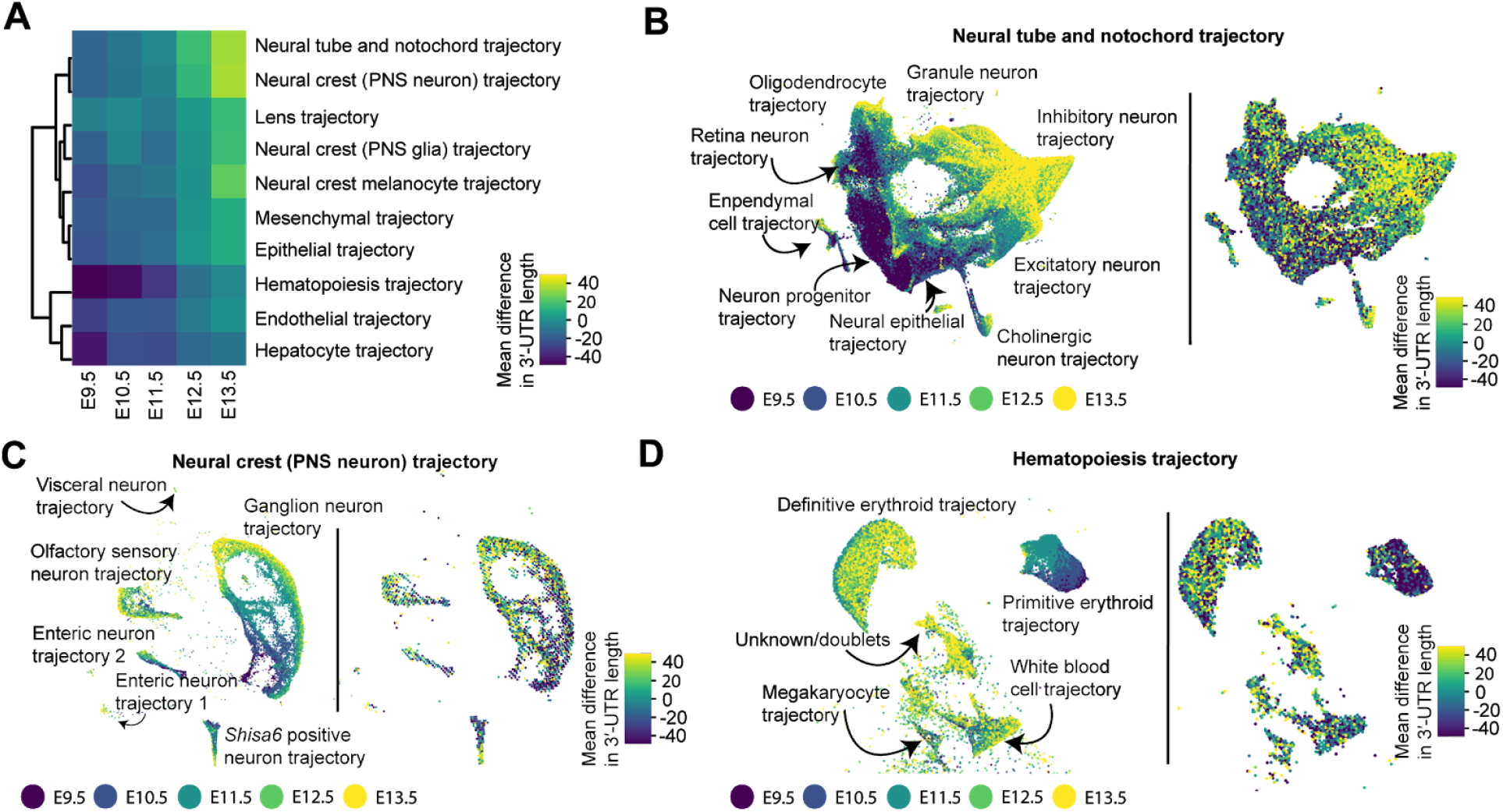
Differential 3′-UTR lengthening among diverse developmental trajectories. **(A)** Heatmap of the mean difference in 3′-UTR length after aggregating cells from each developmental stage and one of ten developmental trajectories computed using UMAP^31^. Heatmap is clustered by Euclidean distance as a distance metric. **(B-D)** UMAP embeddings of the neural tube and notochord (B), neural crest of the peripheral nervous system (C), and hematopoiesis trajectory (D). Cells from each plot are colored by developmental stage (left panel) and mean difference in 3′-UTR length across genes (right panel). Mean differences are calculated with respect to the cells shown in the UMAP rather than all cells.

### Dynamic gene-specific patterns of alternative polyadenylation across early development

While our previous analyses revealed transcriptome-wide trends, it remained unclear how specifically changes in APA manifested at the resolution of individual genes. To investigate this, we tabulated read counts assigned to each gene (*i.e.*, for each PAS and developmental stage, aggregating information across cell types), and used a *χ*^2^ test^30^ to evaluate statistically significant differences in APA for 8,653 genes with at least 100 reads in each of the five developmental stages (**Supplementary Table 2**). This procedure identified 5,169 genes surpassing a False Discovery Rate (FDR) corrected p-value threshold of 0.05. Evaluating the dynamics of the mean 3′-UTR length for this cohort of significant genes at each stage, we discovered that 62% of genes fell into a large cluster that exhibited consistent lengthening over time (**Fig. 4A**). While the majority of these genes showed the greatest increase in lengthening from E11.5 to E12.5, a minority lengthened the most strongly from E10.5 to E11.5 (**Fig. 4A**). About 38% of the significant genes did not simply lengthen across developmental stages, with about half of these progressively shortening over time (**Fig. 4A**). As an alternative method to evaluate transcriptome-wide changes, we computed the entropy across PASs for each gene and developmental stage. This alternative visualization scheme uncovered that nearly 75% of genes obey a progressive decrease in entropy, indicating that as developmental time progresses, a few PASs become increasingly dominant for the vast majority of genes; in contrast, about 15% of genes exhibited the opposite pattern of increased entropy across time, with the remaining displaying heterogeneous patterns (**Supplementary Fig. 5A-B**).

**Figure 4.**
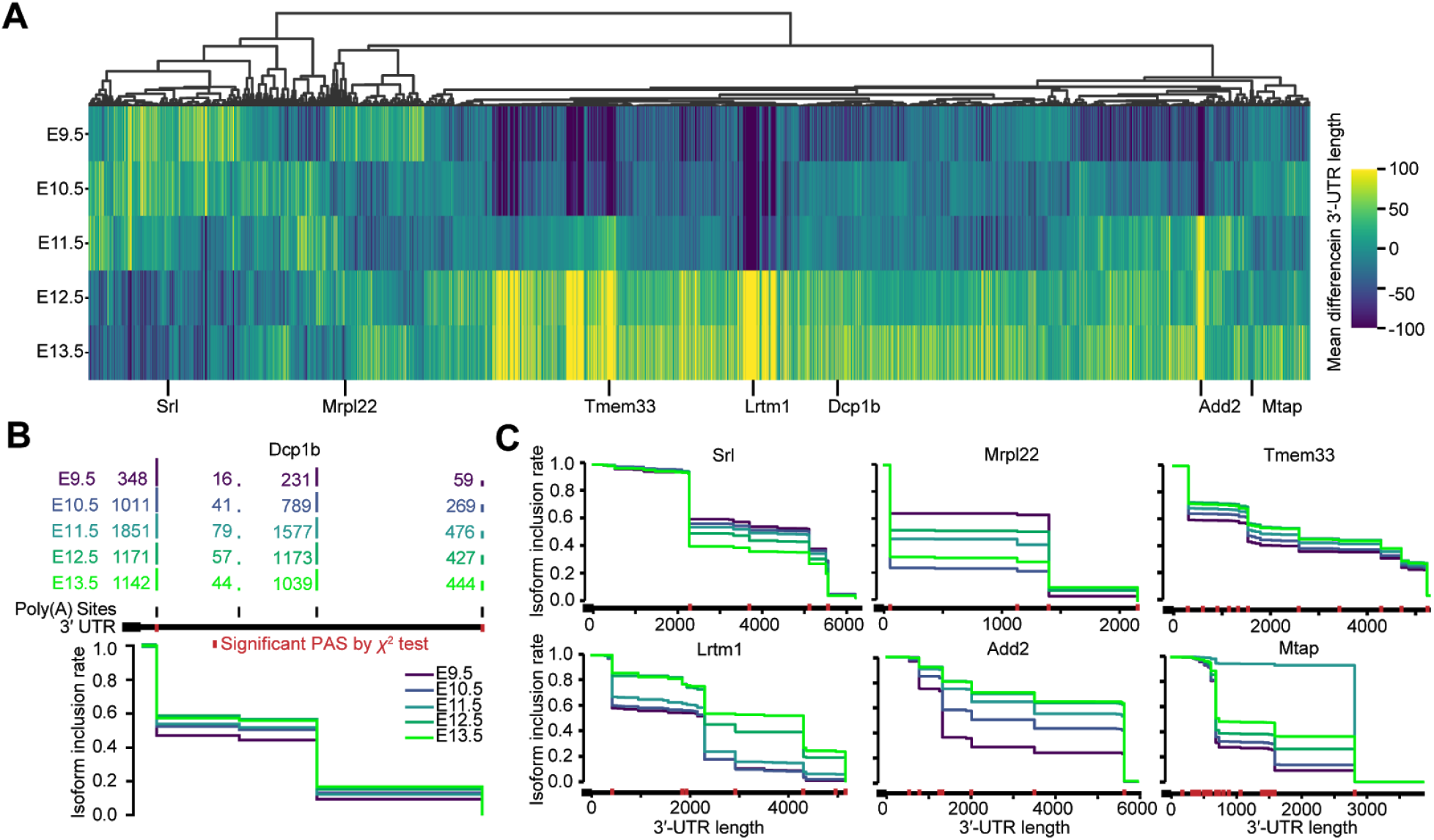
Identification of distinct gene lengthening patterns and genes responsible for overall data trends across embryonic ages. **(A)** Heatmap of mean differences in 3′-UTR lengths for 5,169 genes with significant differences in PAS usage across embryonic stages. Heatmap is column-centered and clustered by Pearson correlation as a distance metric. **(B)** Schematic of IIR plot visualization using PAS counts for each of five embryonic stages. Vertical red lines along the 3′ UTR indicate PASs that are significantly different between stages by the *χ*^2^ test (p<0.05). **(C)** IIR plots for six genes among representative clusters shown in panel (A), and colored by embryonic stage. See also **Supplementary Fig. 5** for clustering according to differences in entropy. See also **Supplementary Table 3** for a table of read counts associated with each PAS for each gene and embryonic stage.

We extended our previous gene-centric IIR plotting scheme (**Fig. 1D**) to visualize the landscape of APA across the five developmental stages assayed, this time using a *χ*^2^ test to highlight individual PASs which were significantly different in at least one stage (**Fig. 4B**). Using this scheme, we visualized an assortment of genes from different clusters to dissect the nature of the isoform switching events contributing to changes in 3′-UTR lengths (**Fig. 4C**). Many of these genes contained dozens of PASs whose relative proportions significantly changed across time. For genes belonging to the dominant cluster (*Tmem33*, *Lrtm1*, *Dcp1b*, and *Add2* in **Fig. 4A**), later developmental stages led to the progressive selection of distal isoforms, leading to progressive 3′-UTR lengthening (**Fig. 4C**). The opposite pattern was observed for a gene belonging to a smaller cluster (*Srl* in **Fig. 4A**), whereby the proximal isoform was selected more frequently over the distal as time progressed, leading to progressive 3′-UTR shortening (**Fig. 4C**). For yet other genes, the choice of distal isoforms was highly time-dependent. For example, *Mtap* displayed a near-complete proximal-to-distal isoform switching event in E11.5, subsequently lengthening beyond baseline levels in later developmental stages; in contrast, *Mrpl22* exhibited progressive shortening, with a dominant distal-to-proximal isoform switching event occurring in E10.5 (**Fig. 4C**).

Finally, we performed a similar gene-centric analysis, this time evaluating differences among individual cell types (*i.e.*, aggregating information across the five developmental stages). Among 1,491 genes with at least 20 reads in each of the 38 t-SNE clusters, we identified 1,078 genes surpassing an FDR-corrected p-value threshold of 0.05, as evaluated by the *χ*^2^ test (**Supplementary Table 4**). This subset of significant genes largely clustered into four cell type groups (C1-C4, **Fig. 5A**) when evaluating differences in mean 3′-UTR length, with cell types within each group displaying strongly correlated patterns across all of the genes. C1, which consisted primarily of neuronal cell types, was unique in that the vast majority of genes displayed global lengthening; conversely, the primitive erythroid lineage was dominated by genes experiencing 3′-UTR shortening (**Fig. 5A**). In special cases, we detected a highly gene-specific and cell-type-specific pattern, as in the case of *Bclaf1* showing 3′-UTR lengthening within a t-SNE cluster annotated as lens cells (**Fig. 5B**). However, for most genes, all of the cell types within each cluster displayed a concerted shift towards either 3′-UTR lengthening (*e.g.*, cluster C1 in *Gnb1*, C1 and C4 in *Samm50*) or 3′-UTR shortening (*e.g.*, cluster C2 in *Polr3k*, C1 in *Hoxd4*) (**Fig. 5B**).

**Figure 5.**
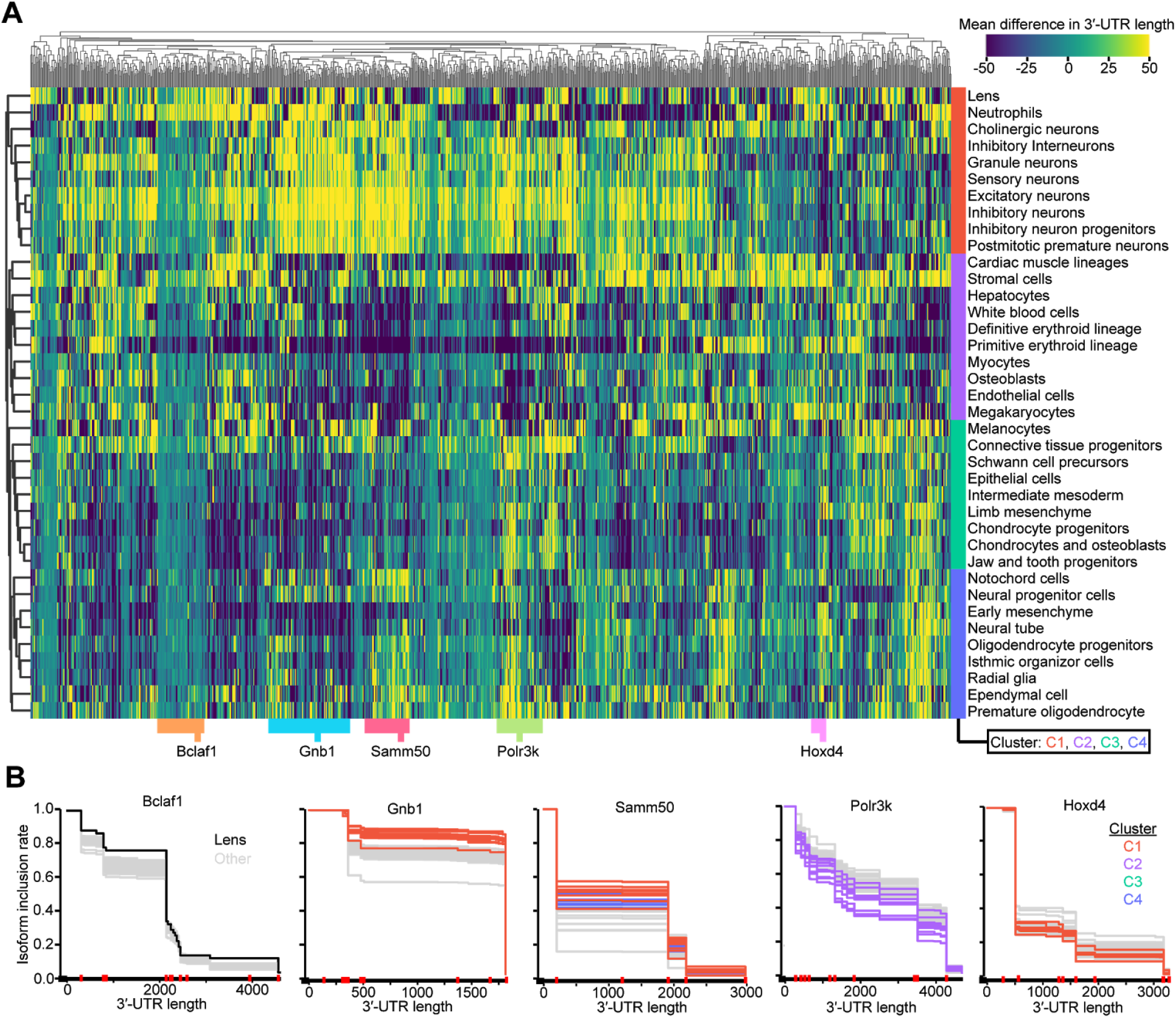
Identification of distinct gene lengthening patterns and genes responsible for overall data trends across embryonic ages. **(A)** Heatmap of mean differences in 3′-UTR lengths for 1,078 genes with significant differences in PAS usage across cell types derived from 38 t-SNE clusters. Heatmap is column-centered and clustered in both rows and columns by Pearson correlation as a distance metric. Row colors indicate the four dominant cell-type clusters. Column colors indicate five representative clusters chosen from the dendrogram above. **(B)** IIR plots for five genes among the representative clusters shown in panel (A), and colored by either individual cell types or cell-type clusters shown in panel (A). Grey lines allude to all other cell types. Vertical red lines along the 3′ UTR indicate PASs that are significantly different between cell types by the *χ*^2^ test (p<0.05). See also **Supplementary Fig. 6** for clustering according to differences in entropy. See also **Supplementary Table 5** for a table of read counts associated with each PAS for each gene and cell type.

When visualizing PAS usage with respect to entropy, several cell types emerged as displaying interesting patterns: the primitive erythroid lineage showed heightened entropy across most genes, whereas neutrophils—and to a smaller degree, the lens—showed decreased entropy (**Supplementary Fig. 6A-B**). This observation suggests a potential for cell-type-specific regulatory mechanisms that guide a more stochastic or more defined choice of PASs, respectively. A smaller subset of genes, such as Sec11a, notably displayed higher entropy among neuronal cell types, consistent with an active mechanism governing a switch towards longer 3′-UTR isoforms (**Supplementary Fig. 6A-B**).

### Putative RNA-binding protein regulators of alternative polyadenylation

Reasoning that changes in the regulation of APA may be coupled to the dynamically changing expression of RNA-binding proteins (RBPs), we searched for RBPs with expression level differences across our five developmental stages and 38 cell types. Having demonstrated that PAS-mapping reads serve as an improved proxy for relative gene expression levels (**Fig. 1E**), we quantified gene expression levels for all protein-coding genes as counts per million (cpm) (**Supplementary Table 6**, **Supplementary Table 7**) and cross-referenced these genes to a database of putative RBPs in the mouse genome^38^.

Evaluating relative differences in RBP expression across our five developmental stages, we observed that about 41% of RBPs exhibited reduced expression across time, while the remaining ones increased or were stable (**Supplementary Fig. 7A**). Performing a similar analysis across our 38 cell types, we observed that about 26% of RBPs were enriched in neuronal lineages (**Fig. 6A**). Interestingly, only a minority of RBPs remained at similar levels in neuronal lineages relative to other cell types, with most being relatively depleted in neurons (**Fig. 6A**). Given our observation that 3′-UTRs lengthen both across the developmental stages and most dramatically in neurons (**Fig. 2C**), we sought to identify putative RBP regulators induced both across time and specifically in neurons relative to other cell types. We therefore quantified the log_2_ fold-change of RBPs in E13.5 relative to E9.5 as well as in neuronal lineages relative to other cell types (**Supplementary Table 8**). We found a strong correlation in RBP expression differences (Pearson *r* = 0.59), with a small group of outliers induced by at least two-fold along both axes (**Fig. 6B**).

**Figure 6.**
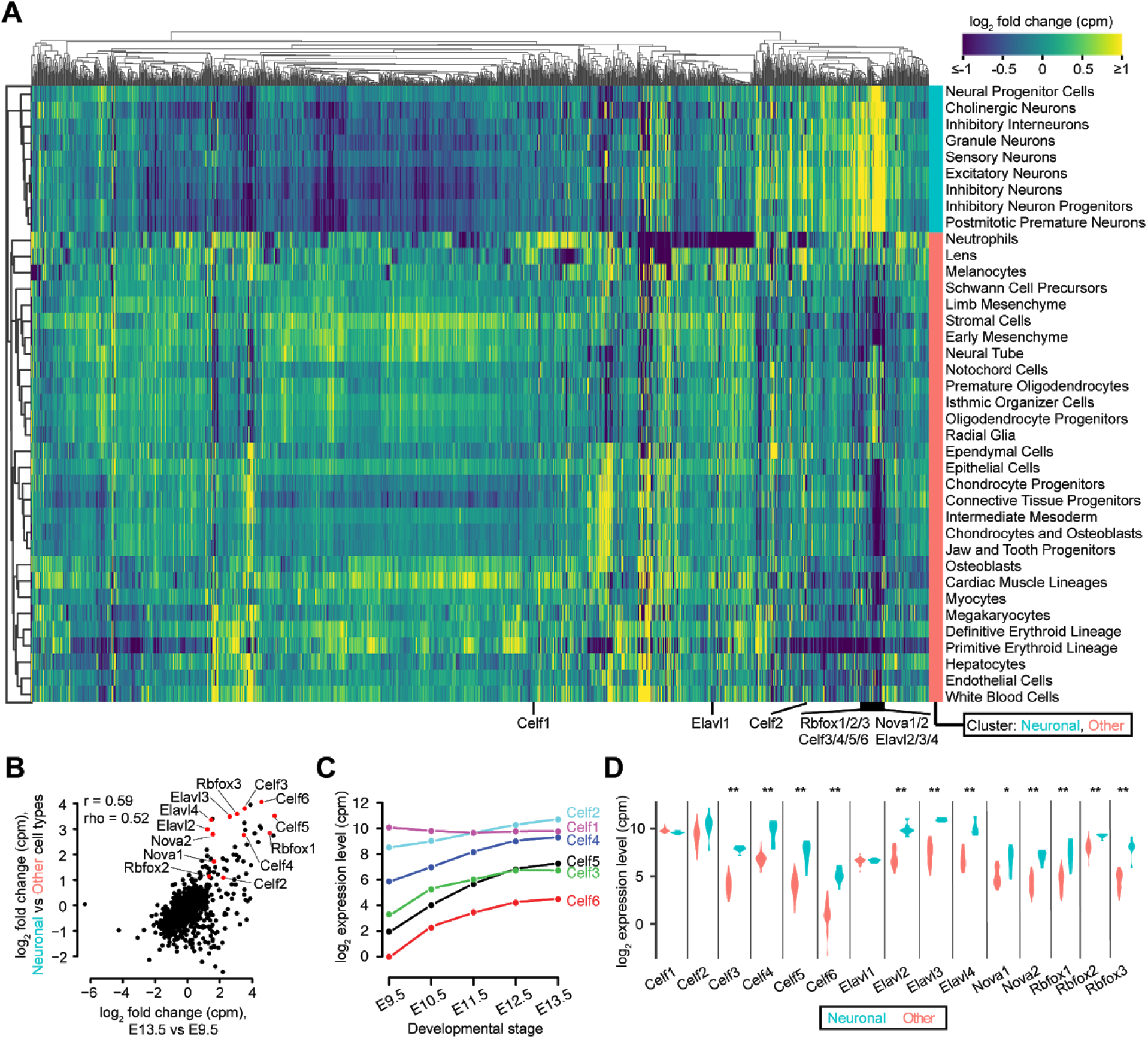
Evaluation of putative RNA-binding protein regulators of alternative polyadenylation. **(A)** Heatmap of relative gene expression levels, quantified as log_2_(counts per million), for a set of 1,576 RBPs across cell types derived from 38 t-SNE clusters. Heatmap is column-centered and clustered in both rows and columns by Pearson correlation as a distance metric. **(C)** Expression levels of *Celf1-6* across the five developmental stages. Expression is quantified in counts per million (cpm) and shown on a log_2_ scale. **(D)** Expression levels of genes highlighted in panel (B) in neuronal cell types relative to other cell types. Significant differences between the two groups were assessed by a two-sided Wilcoxon rank-sum test, with p-values adjusted for multiple hypothesis testing with a Bonferroni correction (*p<0.01, **p<10^-4^).

The most salient factors that were induced comprise a large family of ELAV-like RBPs, including *Elavl2-4* (also known as *HuB*, *HuC*, and *HuD*, respectively) and *Celf2-6* (CUGBP Elav-like family members, also known as *BRUNOL*-*3*, *1*, *4*, *5*, and *6*, respectively) as well as splicing regulators *Nova1-2* and *Rbfox1-3* (**Fig. 6B**). Additional top-ranked RBPs also serve as candidate regulators of APA (**Supplementary Table 8**). The expression of *Elavl2-4*, *Rbfox1-3*, *Celf2-6* monotonically increased in expression across the developmental stages (**Fig. 6C**, **Supplementary Fig. 7B-C**), and were significantly higher in neuronal cell types, while *Elavl1* (also known as *HuR*) and *Celf1* (also known as *BRUNOL*-*2*) remained at similar levels in each context (**Fig. 6D**). These expression patterns are broadly consistent with the known brain-specific and ubiquitous expression associated with ELAV-like^39–41^and Nova^42^ family members, and support the growing functional evidence for *Elavl2-4^43,44^*, *Rbfox2^45^* and *Nova1-2^42^* in the regulation of APA. Additionally, we found that *ELAVL2-4*, *CELF2-6*, and *RBFOX1-3* form an experimentally supported network of protein-protein interactions^46^ (**Supplementary Fig. 7D**).

## DISCUSSION

Despite the rapid growth of single cell RNA sequencing data in recent years, the vast majority of analyses routinely overlook the phenomenon of alternative polyadenylation. Although scRNA-seq was initially developed to measure gene expression levels, multiple orthogonal forms of information are also effectively captured. For example, RNA velocity analysis, which estimates future transcriptome state by modeling intron/exon ratios, illustrates the ability to extract dynamical information about cellular differentiation^47^. In this work, we further develop a computational pipeline to quantify 3′ ends in scRNA-seq data by cross-referencing a novel integrated annotation set of 3′ UTRs and polyadenylation sites, thereby enabling a more spatiotemporally resolved understanding of APA. This pipeline closely recapitulates prior bulk measurements, yet further enables a closer spatiotemporal dissection of APA. Although the utility of scRNA-seq to give insight into APA has been recognized recently^30^, we extend this line of work to an expansive atlas of cell types in a developmental time course spanning multiple stages of embryonic development^31^.

Our findings reinforce the principle that the most proliferative cell types such as blood maintain shorter 3′ UTRs^11,12^, on average, while lowly proliferative ones such as neurons maintain lengthened 3′ UTRs^17,23^. As differentiation progresses, cells of all types naturally become less proliferative, leading to an observed global lengthening of 3′ UTRs in all cell types^22,48^. A major functional consequence of this is that the global shortening of 3′ UTRs could lead to the evasion of microRNA-mediated repression, resulting in greater mRNA stabilities across the transcriptome and enhanced protein synthesis rates in proliferative cells^11^. In contrast to previous work, which often binarized the landscape of 3′ termini into proximal and distal isoforms due to a limited PAS annotation set^22,23^, we develop more general metrics (*e.g.*, changes in mean length and entropy) that consider the relative proportions of the many PASs within each gene. While most genes obey a canonical pattern of 3′-UTR lengthening over time, a small subset of genes deviate from this trend. Most cell types can be grouped into one of four clusters that obey similar trends across genes. These observations are consistent with the evolution of regulatory mechanisms that act in a gene-specific and tissue-dependent manner^1,5^.

An investigation into putative RNA-binding protein regulators that are co-activated in cellular contexts experiencing 3′-UTR lengthening revealed the induction of RBPs of the ELAV-like family, including *Elavl2-4* and *Celf2-6*, as well as splicing regulators *Nova1-2* and *Rbfox1-3*. Prior work provides functional evidence that fly orthologs of the ELAV-like induce neural-specific 3′-UTR lengthening through competition with CstF^43,44,49–51^, and that mammalian *Elavl2-4* can also regulate APA^41,52^. Moreover, *Nova1-2^42^* and *Rbfox2^45^* have been directly implicated as regulators of APA in mouse and rat cells, respectively. Although only *Celf2* has been shown to regulate APA^53^, evidence for the roles of *CELF* family proteins in this process include: i) enriched expression in neurons and later developmental stages, ii) direct interaction with *RBFOX* and *ELAVL* family members, iii) nuclear localization^39^, iv) enriched binding to the 3′-UTR terminus^54^, v) interaction with U2 snRNP^39^, which is known to promote distal isoform usage^55^, and vi) roles in splicing^39,40^.

Given the exaggerated 3′-UTR lengthening we observed in neuronal subtypes, it is interesting to consider how these observations might give insight into regulatory function in neurons. It has been previously observed that APA guides differential mRNA localization^56,57^, and that APA itself is directly regulated by neural activity^58^ such as long-term potentiation^59^. These findings open the possibility that APA might serve as an important process in guiding mRNAs to axons and dendrites, thereby modulating synaptic potential. One promising direction for this work is to use our APA atlas, and those derived from other single cell datasets^60^, to further dissect how differential mRNA localization across neuronal subtypes might contribute to their functional specialization. More generally, we anticipate that our characterization of APA across genes, cell types, and developmental stages of a mammalian organism will serve as a resource to further guide the discovery of new regulatory mechanisms that control APA. It will also help to dissect how these changes impact the function of mRNA with respect to its cellular localization, half-life, and translation in cell types throughout the body.

## METHODS

### An integrated set of mouse 3′ UTRs

We established an integrated set of mouse 3′-UTR annotations for protein-coding genes in which each unique stop codon was associated with a representative transcript with the longest annotated 3′ UTR^9^, using the Gencode M25 “comprehensive” set^33^ as our initial annotations (**Supplementary Table 1**). For each unique stop codon, we selected the longest 3′ UTR from three additional resources: i) RefSeq (March 2020 release)^36^, ii) 3′ UTRs with extreme lengthening^23^, using liftOver^61^ to remap the coordinates from mm9 to mm10, and iii) bulk 3P-seq-based annotations derived from mouse muscle, heart, liver, lung, kidney, brain, testes, and white adipose tissues as well as NIH 3T3 and mESC cell lines^8^. The choose_all_genes_for_TargetScan.pl Perl script in the TargetScanTools Github^35^ was used to integrate these databases, allowing a 3P-seq read to exist up to 5,400nt (*i.e.*, the 99th percentile of annotated 3′-UTR lengths) downstream of a stop codon.

In certain scenarios, such as in the case of alternative splicing of the terminal exon, a gene is potentially associated with many unique stop codons, each of which is associated with its own 3′-UTR annotation. We therefore sought to avoid a bias in which genes with many such transcript isoforms would be overrepresented in the downstream results, and to avoid the possibility that PASs would be counted redundantly in cases in which multiple different 3′ UTRs overlapped the same genomic coordinates. We therefore carried forward a single transcript isoform with the greatest number of 3′-UTR mapping reads (or a random top-ranked one in the case of a tie) to represent each gene. To perform this counting, scRNA-seq reads were intersected with our 3′-UTR annotation set using bedtools intersect (-wa -wb -s)^62^.

### An integrated set of mouse poly(A) sites

To generate our union PAS set, we integrated three PAS annotation databases: Gencode M25^33^, PolyA_DB v3^20^, and PolyASite 2.0^21^. First, PASs within ±10nt of another PAS within the same database were collapsed by selecting the most downstream PAS. Next, the following procedure was implemented to reduce redundancy between databases: i) we collected PASs from PolyASite 2.0, ii) we added PASs from PolyA_DB v3 not within ±10nt of the current PAS set, and iii) we added PASs from Gencode M25 not within ±10nt of the current PAS set. This method of sequential addition led to a total of 164,772 PASs in our union set; we provide the genomic coordinates and corresponding read counts associated with this set (**Supplementary Table 3**, **Supplementary Table 5**).

### Calculation of 3′-UTR lengths, relative length differences, and corresponding visualizations

Reads were mapped to the mm10 genome and collected from previous work^31^ (GEO ID: GSE119945). 3′-UTR length corresponding to a given read was computed as the distance from the stop codon to the read’s assigned PAS, minus the length of any intervening intron(s). These 3′-UTR lengths were used to compute a “gene by cell” sparse matrix of the mean length among all 3′-UTR isoforms, weighted by their respective counts. For each gene, we then computed each cell’s deviation from the mean of 3′-UTR lengths across cells, considering only non-missing values. For heatmaps, these deviation values were then averaged according to the labels assigned to each cell (*i.e.*, with respect to t-SNE cluster, UMAP trajectory, and/or developmental stage). Cell labels were based upon those previously assigned^31^. When indicated in the legend, in some instances the heatmaps were further centered by subtracting the mean of the row or column. t-SNE plots were visualized using the hexbin (gridsize = 500, vmin = -50, vmax = 50) function from pyplot, which averages values from cells captured in local bins. UMAP plots were binned by splitting each of the x, y, and z coordinates into 150 equally sized bins.

### Gene-level isoform inclusion rate plots and corresponding statistics

For each gene, we counted reads assigned to each PAS to build contingency tables of counts for either (PAS by developmental stage) (**Supplementary Table 3**) or (PAS by cell type) (**Supplementary Table 5**). We then computed statistical significance using the *χ*^2^ test as computed by the chisquare function in scipy, either with respect to the entire gene (**Supplementary Table 2, Supplementary Table 4**) or with respect to each PAS (axis = None or axis = 1, respectively). In both cases, we provided a matrix of expected counts, based on the joint probability of each cell multiplied with the total counts in the matrix. For the gene-level *χ*^2^ test, we further derived a Benjamini-Hochberg (BH) based q-value to account for the FDR. Considering the read counts associated with each PAS position, isoform inclusion rates were visualized in the same manner as previous work, which allude to this plotting style as the Affected Isoform Ratio (AIR) plot^8,9,35^. Much like a survival curve, the IIR quantifies the proportion of 3′-UTR isoforms that include a given nucleotide position.

### Search for putative RBP regulators

To evaluate changes in gene expression associated with RBPs, we first computed gene expression levels for all protein-coding genes. Towards this goal, we summed the counts associated with PAS-mapping reads for all unique PASs (*i.e.*, as assessed by chromosomal coordinate) across all transcripts (*i.e.*, including those with alternative last exons) corresponding to each gene, using our count tables partitioned either by developmental stage (**Supplementary Table 3**) or by cell type (**Supplementary Table 5**). Counts were then normalized by the stage or cell type into counts per million (cpm) (**Supplementary Table 6, Supplementary Table 7**) and then log_2_-transformed. Genes were annotated as an RBP if their gene name matched one of 1,882 mouse genes annotated as a putative RBP^38^. For the subset of 1,576 RBPs meeting an expression threshold of 2 cpm in at least one of the samples tested, we computed the fold-change of each gene across time as [log_2_(cpm at E13.5) – log_2_(cpm at E9.5)] and in neurons relative to other cell types as [mean log_2_(cpm in neurons) – mean log_2_(cpm in other cell types)] (**Supplementary Table 8**), where neurons are defined as the cell types in the cluster shown in **Fig. 6A**.

## AUTHOR CONTRIBUTIONS

V.A. conceived of the study, designed analyses, and performed RBP analysis. S.L-D. performed the remaining computational analyses and generated tables. V.A. and S.L-D. generated figures and wrote the paper with feedback from D.R.K. and J.S.

## ACKNOWLEDGEMENTS

We thank Junyue Cao and other members of the Shendure Lab for discussions surrounding the single cell data as well as Nimrod Rubenstein for critical commentary. We are grateful to Eric Lai for discussions surrounding RBP analysis and ELAV-related regulatory mechanisms. This material is based upon work supported under an NRSA NIH fellowship 5T32HL007093 (to V.A.). J.S. is an investigator of the Howard Hughes Medical Institute.

## SUPPLEMENTAL TABLES

**Supplementary Table 1.** GTF file of the gene models associated with the “Integrated 3′ UTR” set, as described in **Fig. 1D**, along with the source database of the 3′-UTR annotation. Genomic coordinates are provided with respect to the mm10 genome.

**Supplementary Table 2.** Table of FDR-corrected q-values for each transcript/gene tested for differential PAS usage across the five developmental stages. Genes that did not pass the threshold of having 100 reads at each age are listed as “Not Tested”.

**Supplementary Table 3.** Table of read counts associated with each PAS for each gene and developmental stage. Genomic coordinates are provided for every PAS with respect to the mm10 genome, along with the annotation source of the PAS.

**Supplementary Table 4.** Table of FDR-corrected q-values for each transcript/gene tested for differential PAS usage across the 38 cell types. Genes that did not pass the threshold of having 20 reads in each cell type are listed as “Not Tested”.

**Supplementary Table 5.** Table of read counts associated with each PAS for each gene and cell type. Genomic coordinates are provided for every PAS with respect to the mm10 genome, along with the annotation source of the PAS.

**Supplementary Table 6.** Table of expression levels of protein-coding genes, computed as counts per million, across the five developmental stages. Genes are annotated according to which correspond to known RNA binding proteins.

**Supplementary Table 7.** Table of expression levels of protein-coding genes, computed as counts per million, across the 38 cell types. Genes are annotated according to which correspond to known RNA binding proteins.

**Supplementary Table 8.** Table of log_2_(fold changes) of expression levels for RBPs tested in E13.5 vs E9.5 and in neuronal lineages vs other cell types, ranked by their degree of upregulation in neurons.

## SUPPLEMENTAL FIGURES

**Supplementary Figure 1.**
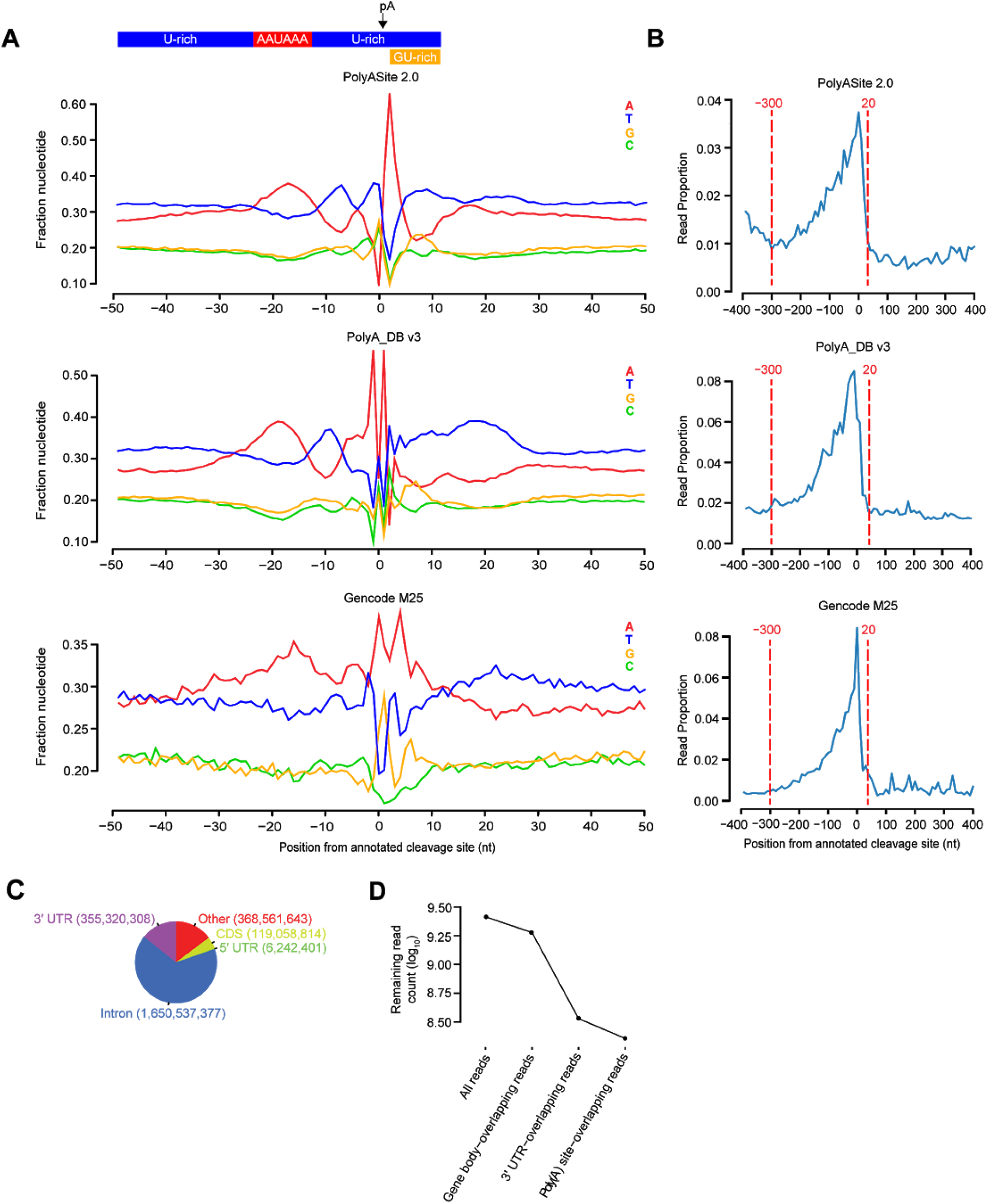
Characterization of PAS annotation databases and properties of scRNA-seq data read filtering. **(A)** This panel is the same as that shown in **Fig. 1B**, except displays information for cleavage sites anchored on PASs unique to each of the three databases (*i.e.*, non-intersecting with a PAS from any other database). **(B)** This panel is the same as that shown in **Fig. 1C**, except displays information for cleavage sites anchored on PASs unique to each of the three databases (*i.e.*, non-intersecting with a PAS from any other database). **(C)** Pie chart showing the relative proportions of scRNA-seq reads that map to each functional region within the genome. **(D)** Plot of the decay in the numbers of reads remaining after each sequential filtering step that was required to isolate the subset of PAS-mapping reads.

**Supplementary Figure 2.**
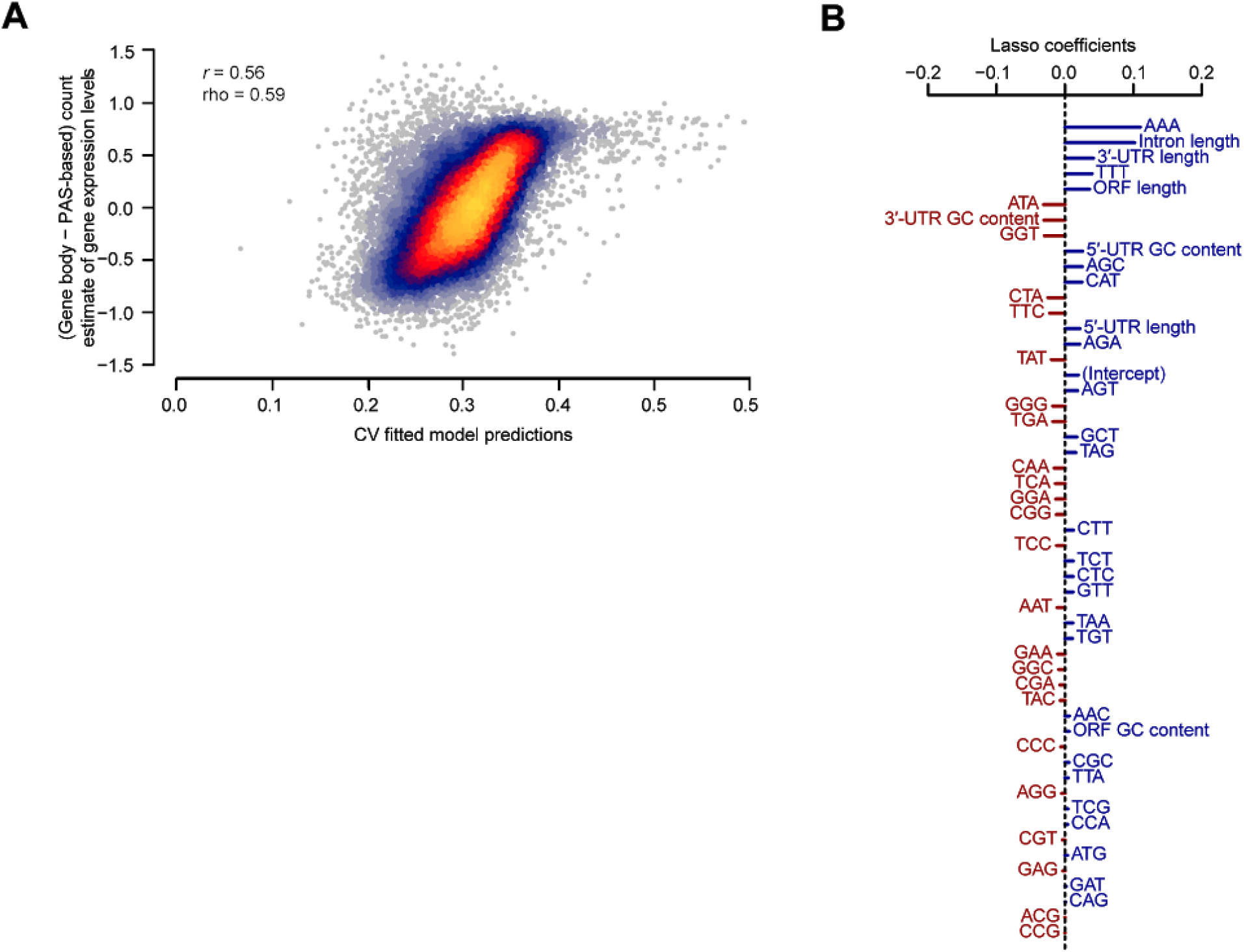
Sequence-based features partially explain biases in the gene-body-based method of estimating relative gene expression levels. We developed a regression model to predict the difference between the gene body and PAS counting methods to estimate gene expression levels, as shown on the x-axes of the left and right panels, respectively, of **Fig. 1E**. The features considered in the model include the length of the 5′ UTR, ORF, introns, and 3′ UTR; the GC content of the 5′ UTR, ORF, and 3′ UTR; and the proportions of each of 64 possible 3-mers within the entire gene body. All features were z-score transformed to enable comparisons between regression coefficients. Following our previous work^63^, we trained a lasso regression model using these features. The strength of the regularization was controlled by a single λ parameter, which was optimized using 10-fold cross-validation for each training set using the *cv.glmnet* function of the *glmnet* library in R. **A)** Scatter plot displaying the relationship between the 10-fold cross-validated predictions derived from the lasso regression model and the observed difference between gene body and PAS-based counting methods of estimating gene expression level. Also indicated are the Pearson (*r*) and Spearman (rho) correlation values. **B)** The ranked coefficients derived from a lasso regression model trained on the full dataset. Positive coefficients (blue) are associated with inflated gene body read counts, while negative coefficients (red) are associated with underrepresented gene body read counts.

**Supplementary Figure 3.**
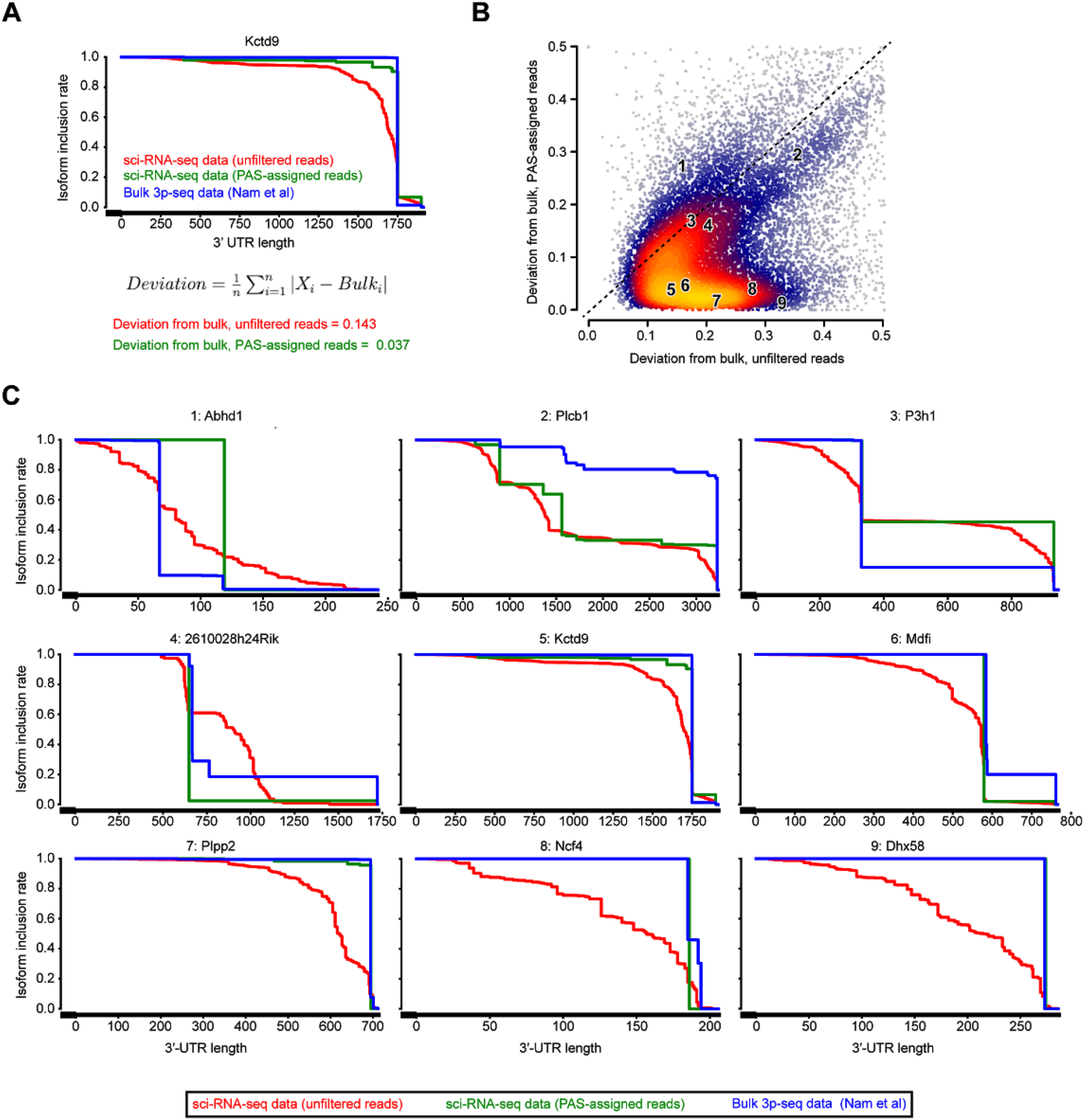
Improvement in the quantification of 3′-UTR isoform abundances after read-to-PAS assignment. **(A)** Calculation of the Mean Absolute Deviation (MAD) metric on an example gene to quantify the divergence between IIR profiles derived from either raw scRNA-seq data or post-processed data after read-to-PAS assignment, relative to the profile for bulk 3P-seq data^8^ as a gold standard. Larger numbers indicate poorer agreement. **(B)** Scatter plot of MAD values for either raw scRNA-seq data (x-axis) or post-processed data after read-to-PAS assignment (y-axis) (n = 16,334 protein-coding genes). Regions are colored according to the density of data from light blue (low density) to yellow (high density). 78% of genes exist below the diagonal dotted line, indicating an improved similarity to bulk measurements after post-processing. Nine genes from different representative regions of the plot are indicated. **(C)** IIR plots for the nine representative genes numbered in panel (B). The majority of genes show strongly improved agreement to bulk 3P-seq data (genes 5-9).

**Supplementary Figure 4.**
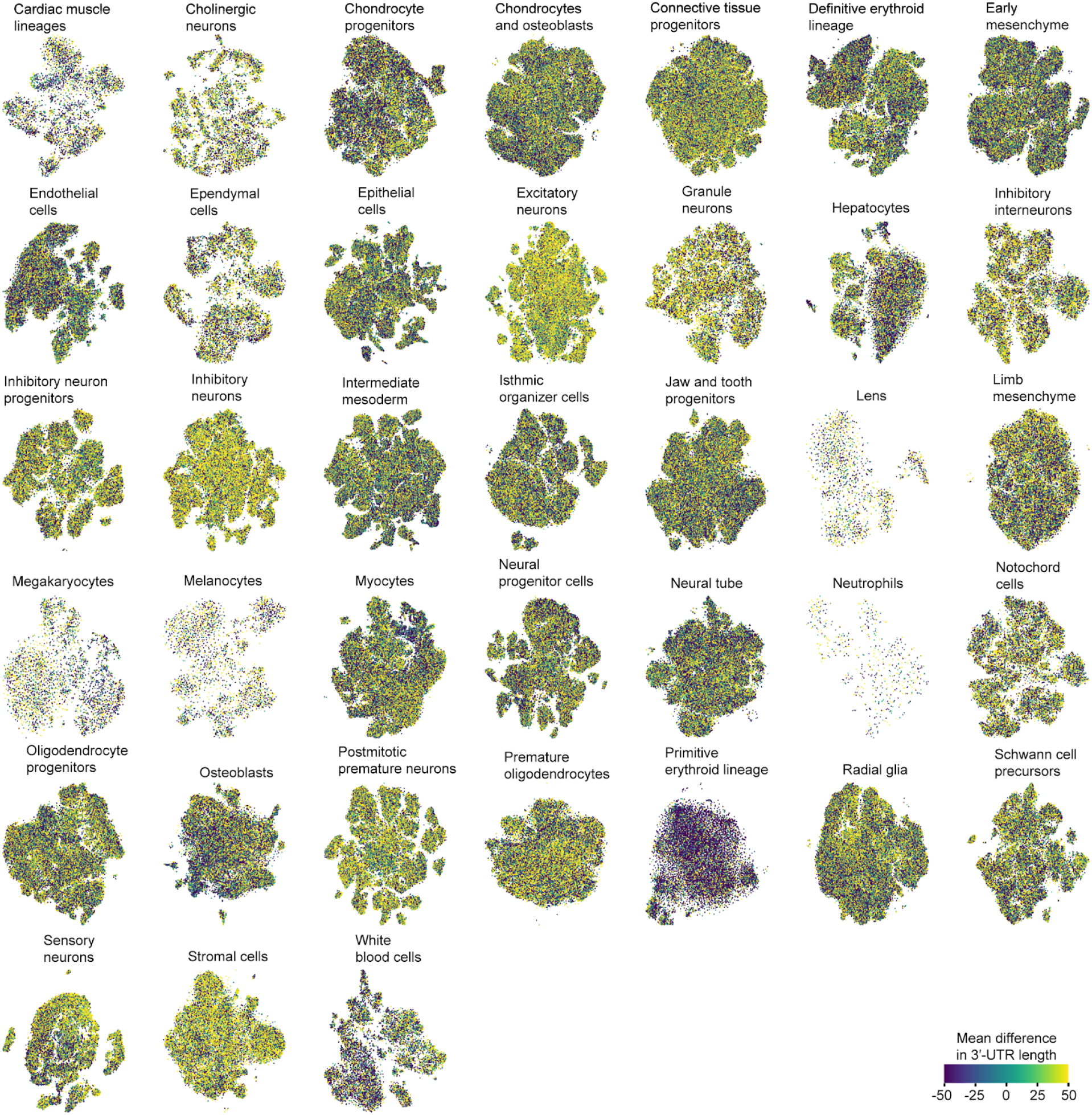
Analysis of differential 3′-UTR lengthening among cellular subtypes. t-SNE embeddings were generated to identify cellular subtypes for cell types derived from each of 38 t-SNE clusters^31^. Each cell was colored according to the mean difference in 3′-UTR lengths across all genes. Local subcluster heterogeneity can be observed (*e.g.*, cellular subtypes in hepatocytes and osteoblasts), along with global differences between clusters (*e.g.*, the primitive erythroid lineage relative to neuronal cells and stromal cells).

**Supplementary Figure 5.**
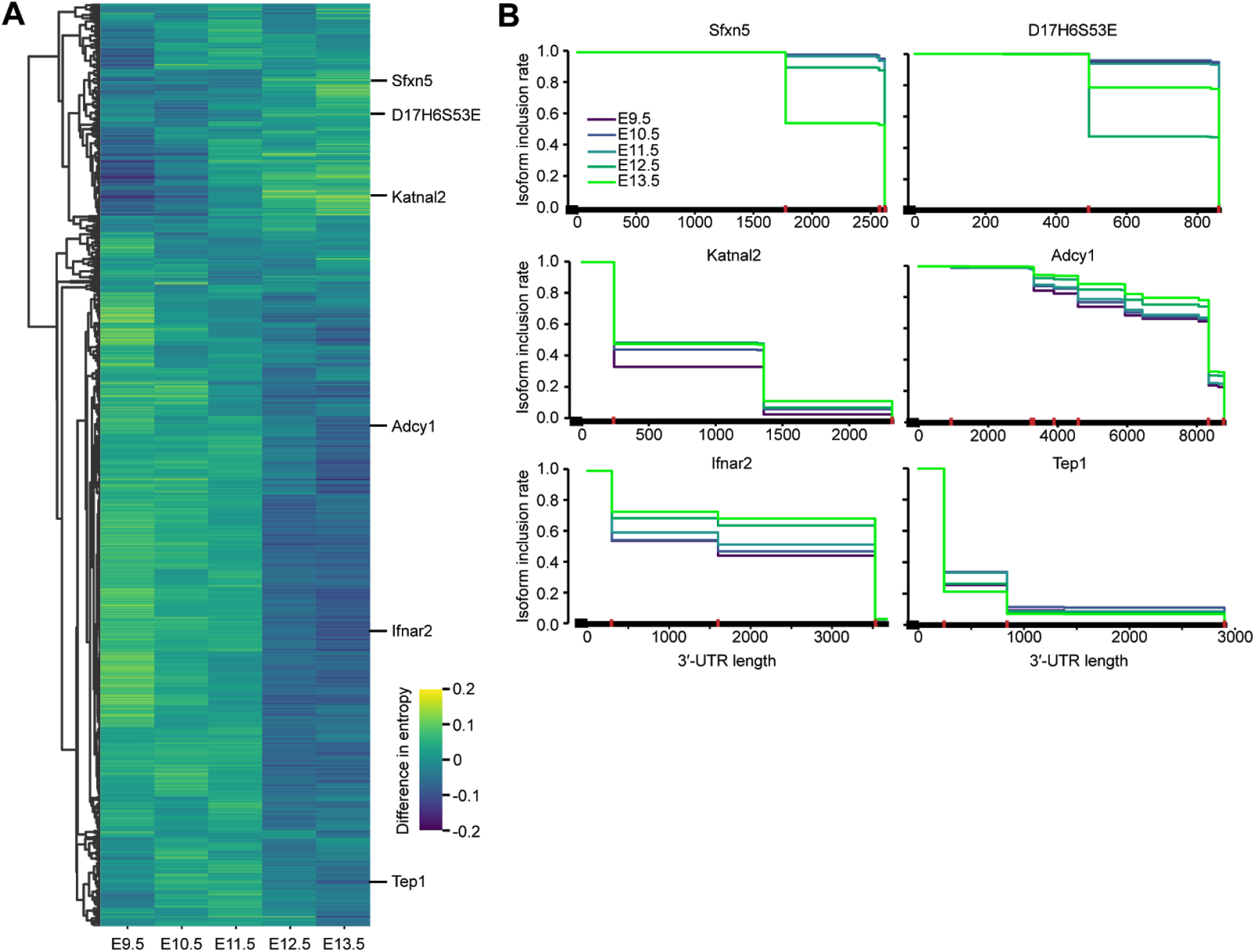
Evaluation of differential PAS usage and diversity across developmental time. **(A)** Heatmap of differences in entropy for the statistically significant genes from panel (A). Entropy for a given gene and developmental stage was calculated as 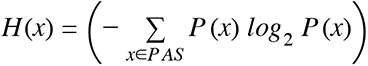 using relative read proportions assigned to each PAS for the set of all PASs associated with the gene. Heatmap is row-centered and clustered by Pearson correlation as a distance metric. Higher values of entropy represent a greater degree of randomness and uniformity in selection among multiple PASs; conversely, lower values represent greater fidelity in selection amongst fewer PASs. **(B)** IIR plots for six genes among representative clusters shown in panel (B), and colored by developmental stage. Vertical red lines along the 3′ UTR indicate PASs that are significantly different between stages by the *χ*^2^ test (p<0.05).

**Supplementary Figure 6.**
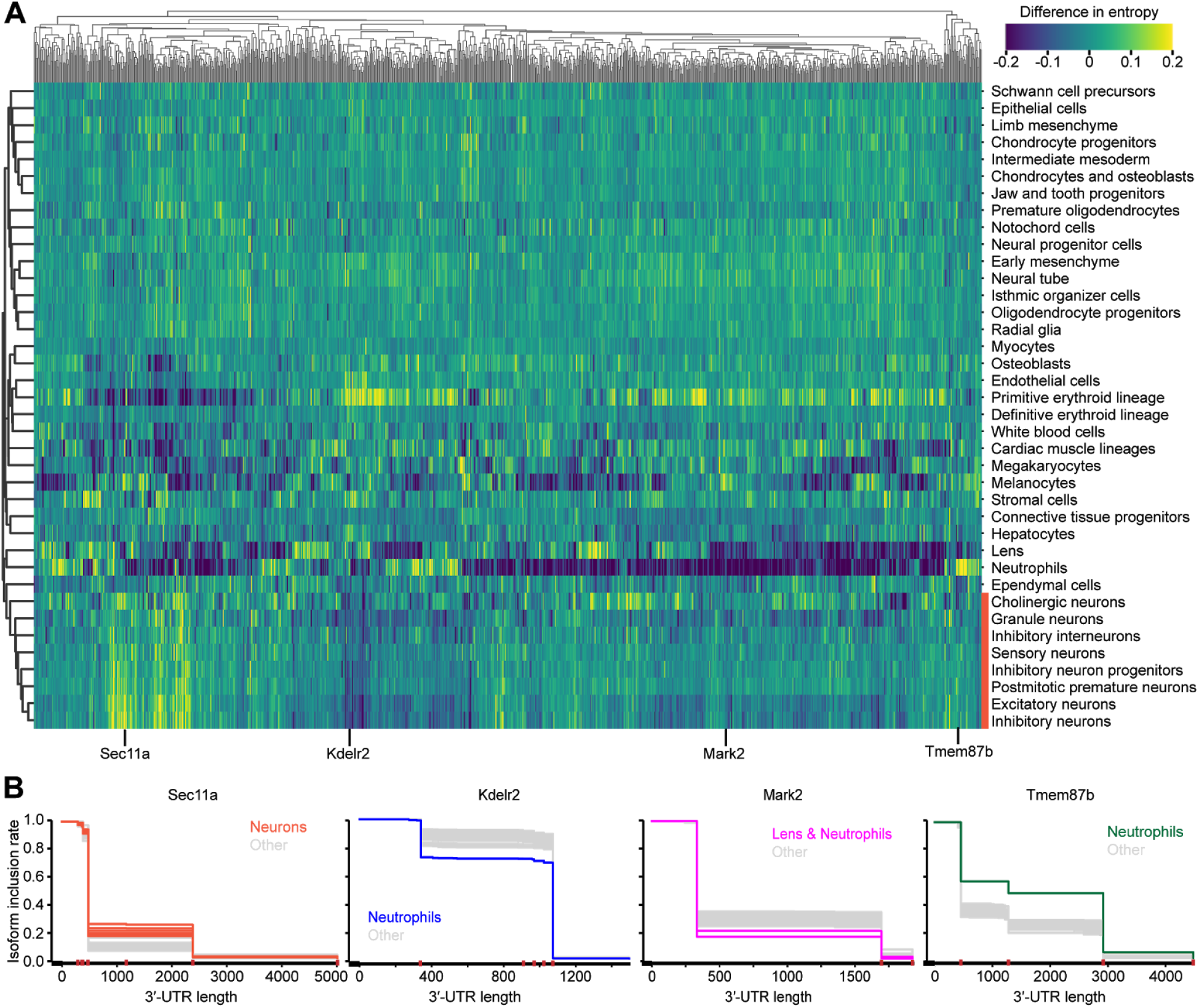
Evaluation of differential PAS usage and diversity across cell types. **(A)** Heatmap of differences in entropy for the statistically significant genes from panel (A). Entropy for a given gene and cell type (derived from each of 38 t-SNE clusters) was calculated as 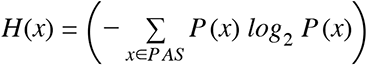 using relative read proportions assigned to each PAS for the set of all PASs associated with the gene. Heatmap is column-centered and clustered in both rows and columns by Pearson correlation as a distance metric. Higher values of entropy represent a greater degree of randomness and uniformity in selection among multiple PASs; conversely, lower values represent greater fidelity in selection amongst fewer PASs. **(B)** IIR plots for five genes among representative clusters shown in panel (B), and colored either by the indicated cell types or the cluster of neuronal cell types shown in panel (B). Grey lines allude to all other cell types. Vertical red lines along the 3′ UTR indicate PASs that are significantly different between cell types by the *χ*^2^ test (p<0.05).

**Supplementary Figure 7.**
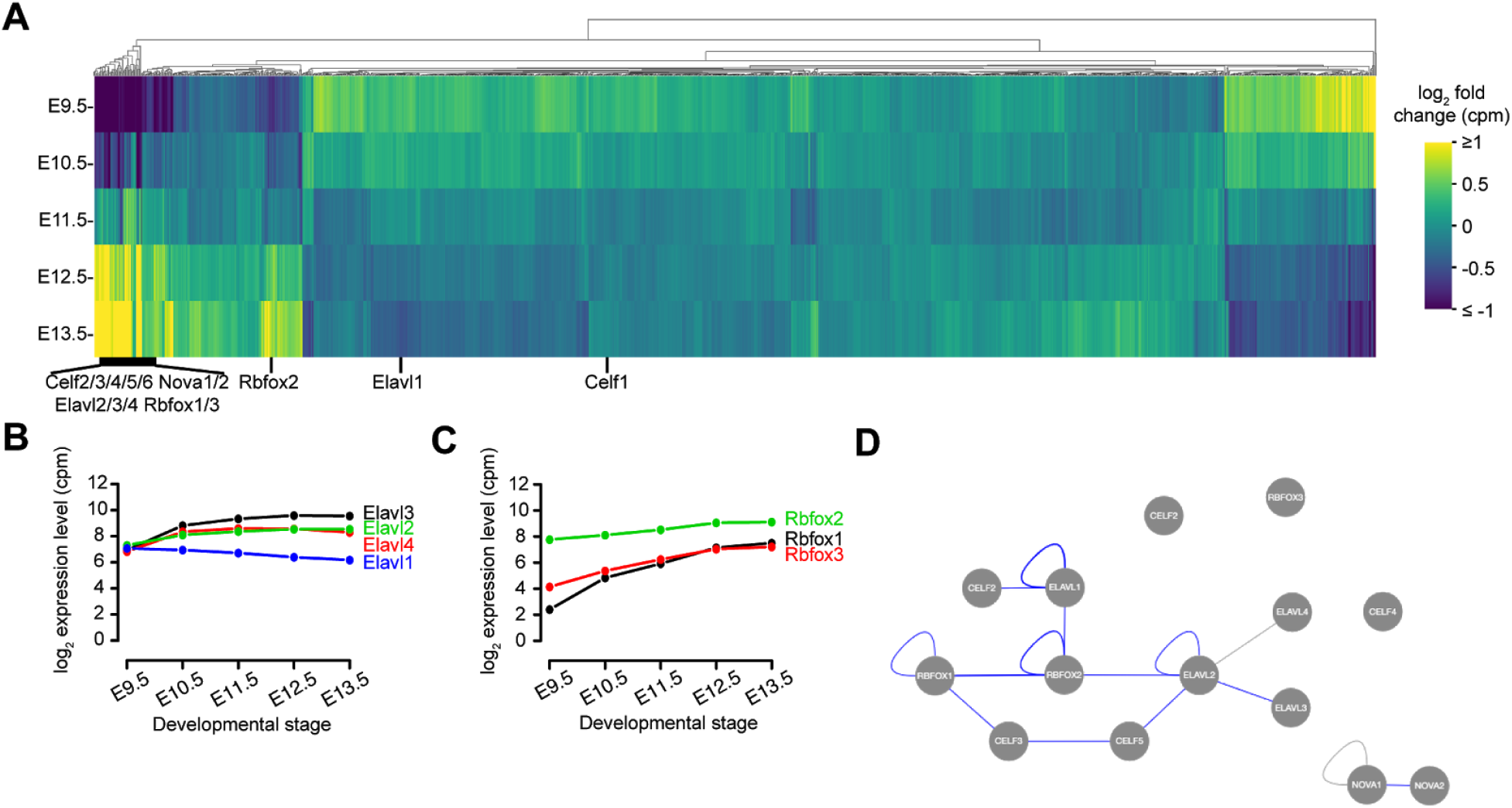
Evaluation of differential RNA-binding protein expression across developmental time. **(A)** Heatmap of relative gene expression levels, quantified as log_2_(counts per million), for a set of 1,516 RBPs across the five embryonic stages. Heatmap is column-centered and clustered by Euclidean distance as a distance metric. (**B**-**C**) Expression levels of *Elavl1-4* (B) and *Rbfox1-3* (C) across the five developmental stages. Expression is quantified in counts per million (cpm) and shown on a log_2_ scale. **(D)** Network of protein-protein interactions supporting RBP interactions as observed in the APID database^46^. Blue lines indicate additional experimental support for the interaction.

